# Energy-ordered resource stratification as an agnostic signature of life

**DOI:** 10.1101/2024.03.27.586986

**Authors:** Akshit Goyal, Mikhail Tikhonov

## Abstract

The search for extraterrestrial life hinges on identifying biosignatures, often focusing on gaseous metabolic byproducts as indicators. However, most such biosignatures require assuming specific metabolic processes. It is widely recognized that life on other planets may not resemble that of Earth, but identifying biosignatures “agnostic” to such assumptions has remained a challenge. Here, we propose a novel approach by considering the generic outcome of life: the formation of competing ecosystems. We use a minimal model to argue that the presence of ecosystem-level dynamics, characterized by ecological interactions and resource competition, may yield biosignatures independent of specific metabolic activities. Specifically, we propose the emergent stratification of chemical resources in order of decreasing energy content as a candidate new biosignature. While likely inaccessible to remote sensing, this signature could be relevant for sample return missions, or for detection of ancient signatures of life on Earth itself.

---

The search for extraterrestrial life hinges on the identification of biosignatures, markers indicating the presence of biotic processes [1]. By necessity, all proposed candidates are inspired by the kind of life we know — the forms of life as they exist on Earth. The type of biosignature receiving the most attention is the expected presence of certain gaseous metabolic byproducts [2, 3], which could be detected spectroscopically, with much attention devoted to specifically molecular oxygen [4]. This kind of signatures assumes that the biotic processes we seek to detect rely on specific chemical transformations for their metabolism.

It is widely recognized that life elsewhere in the universe need not resemble the terrestrial form, motivating the interest in so-called agnostic biosignatures, those that are “not tied to a particular metabolism-informational biopolymer or other characteristic of life as we know it” [5–7]. Some proposals include looking for polyelectrolytes [7] and homochirality [8], but identifying agnostic biosignatures is a serious challenge. In fact, it is unclear if truly agnostic signatures can exist without a theory of life [9]. It is fair to say that all proposed signatures require some additional assumptions, e.g., metabolism, morphology, chirality. Even setting aside the technological challenges of remote sensing, and assuming we could perform arbitrary measurements (e.g., to detect ancient signatures of life on Earth, which is also an active field of research), it is not clear what to look for without making such assumptions. The one thing we all agree on is that life requires self-replication supporting a Darwinian process. However, no measurable signature is known to be a generic consequence of self-replication alone.

Here, we focus on the observation that, as far as we know, life forms never exist alone, but generically develop into systems characterized by ecological interactions and resource competition (even in experimental conditions specifically intended to avoid this [10]). On Earth, there is only one known exception [11]. This suggests that the assumption of life forming an ecosystem is almost as weak as the minimal required assumption of a Darwinian process. This motivates us to ask: are there any biosignatures one might expect to generically arise from the fact that life results in a competing ecology?

We stress that even biosignatures associated with specific metabolisms are generally assumed to be part of ecosystem-level processes. For instance, isotopic fractionation of sulfur, discussed as a candidate biosignature [12], effectively requires an ecosystem-level sulfur cycle. In this sense, many signatures of life are already understood to be signatures of ecosystems. However, the biosignatures usually considered are imprinted by some specific metabolic process this ecosystem is assumed to run. In contrast, here we ask whether there are signatures expected to arise from the competitive nature of ecosystem dynamics, rather than specific metabolic activities.

We propose that one such signature is the emergent stratification of chemical resources in order of their energy content (Fig. 1). Such patterns are observed in many contexts on Earth [13– Abiotically, there is no reason to expect this, as the abiotic rate and energy yield of a chemical reaction are set by independent parameters (height of the activation barrier, *vs*. energy of final state), and are generically uncorrelated. Using a minimal theoretical model, we demonstrate that energy-ordered stratification is a robust consequence of two processes: biological self-replication as species consume resources, and ecological interactions between different biological species as they compete for space. Our model does not assume any specific molecular detail or metabolism. Thus, we propose energy-ordered resource stratification as a candidate agnostic biosignature requiring minimal assumptions on the chemical implementation of the Darwinian process.

**FIG. 1.**
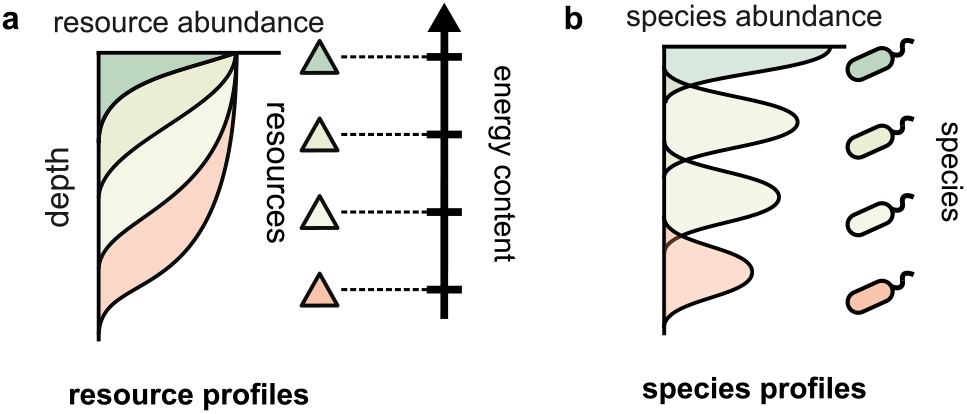
Energy-ordered resource stratification as an observable signature of life. (a) Stratified profiles of chemical resources layered by energy content are commonly observed on Earth, e.g., microbial mats, early Earth fossils (stromatolites), Winogradsky columns, and in marine environments. (b) Such profiles are generally understood to be shaped by biotic species (typically microbes) that metabolize these resources for energy. Here, we propose that energy-ordered resource stratification is a robust signature of biotic action.

## THE MODEL

We seek to understand the consequences of self-replication and ecological interactions for biosignatures. To this end, we consider the following minimal model implementing these two ingredients (Fig. 2a) in a simple setting. We track the dynamics of the abundance of *S* reaction catalysts *N*_*i*_(*x, t*) and *M* chemical resources *R*_*α*_(*x, t*) along a one-dimensional spatial coordinate *x* (representing, e.g., depth in a microbial mat or water column). All reaction catalysts consume different resources and diffuse over space with diffusion constant *D*_*N*_ . A global parameter *γ* controls whether reaction catalysts are biotic and can self-replicate: *γ* = 0 corresponds to abiotic catalysis, while *γ≠* 0 corresponds to self-replicating biological species, which grow as they consume resources. To maintain the growth of these species, resources are supplied abiotically from the outside at *x* = 0 at a constant flux *K*_*α*_. Thus, over time, resources are supplied, depleted and diffuse over space with diffusion constant *D*_*R*_. For a simple implementation of ecology, we assume that biological species at the same location *x* also compete with each other for space with a pairwise competitive interaction strength *α*_*ij*_*N*_*i*_(*x, t*)*N*_*j*_(*x, t*). A global parameter *ρ* controls the density of ecological interactions by controlling the fraction of non-zero entries in the matrix *α*_*ij*_; *ρ* = 0 corresponds to the case of no ecology. In this manuscript, we will only tune the key parameters *γ* and *ρ*, representing self-replication and ecology, while keeping the rest fixed. Finally, we assume for simplicity that species and resources cannot leave the system, while resources enter at *x* = 0 at flux *K*. With these assumptions, the equations governing the dynamics of our model can be expressed as follows:

**FIG. 2.**
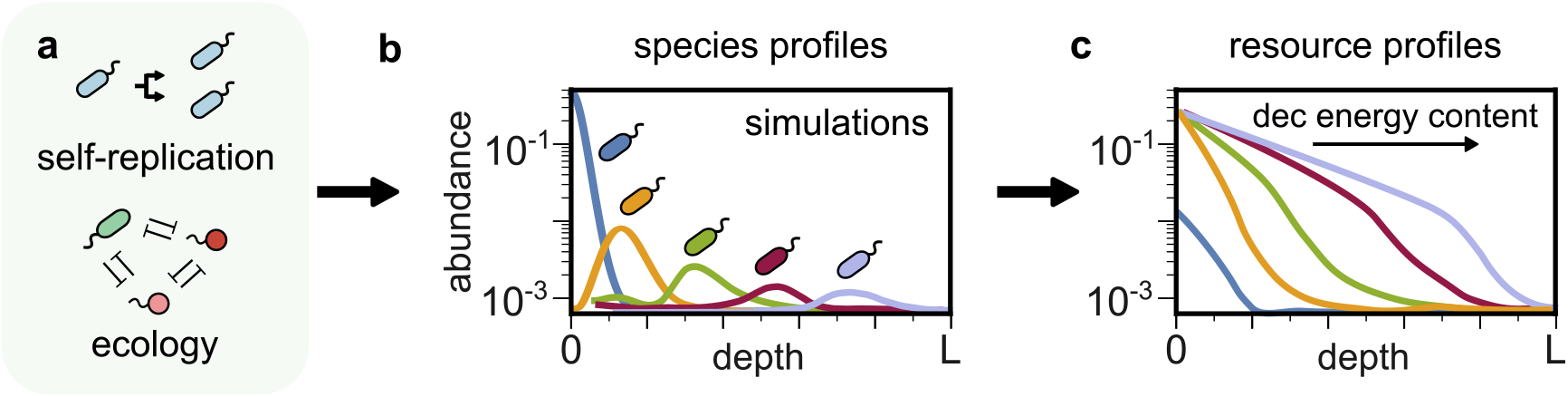
Self-replication and ecology lead to energy-ordered resource stratification. (a) Two universal features of life are self-replication and ecological interactions between different biological species — the simplest being antagonism. (b) Simulating a minimal model incorporating these two ingredients (for details see text) show these two ingredients lead to spatially stratified profiles of (b) species and (c) resources. Shown here is an example from a simulation for 5 species and 5 resources. Antagonistic interactions segregate species spatially, with species displacement order determined by the energy content of the resource they consume. In each segregated zone, species deplete resources proportional to their abundance.

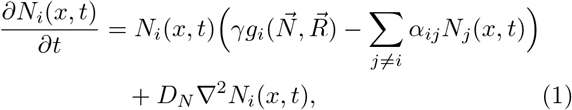

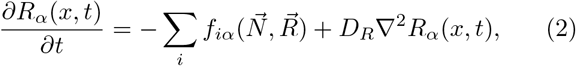

where the self-replication parameter *γ* controls the degree of self-replication (*γ* = 0 meaning no self-replication). The growth of biological species *g*_*i*_ and consumption of resources *f*_*iα*_ have the following functional forms:

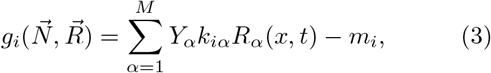

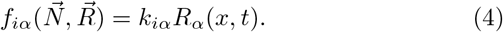

The parameter *Y*_*α*_ represents the energy content of resource *α* and *m*_*i*_ represents the maintenance energy for species *i*. For simplicity, we assume that each biological species is a specialist and consumes only one resource.

If the system were well-mixed, the single “fittest” biological species using the most energy-dense resource would outcompete all others and take over the system. In our model, diffusion across space promotes the coexistence of species using resources with different energy content.

## RESULTS

Simulations of our model show that self-replication and ecology lead to spatial stratification of biological species and chemical resources (Fig. 2b–c). Starting from no stratification, with homogeneous species profiles and identical resource profiles, species and resource dynamics naturally converge towards a steady state where the biological species, and consequently resources, become stratified (Fig. 2b–c). The emergence of stratification can be understood as follows: the species using the most energetic resource grows the fastest near the source *x* = 0. Because it grows fastest, this species most strongly antagonizes all others near *x* = 0 (Fig. 2b, blue). As a result, less energetic resources are not consumed near *x* = 0 and penetrate further (Fig. 2b, orange, green, red and purple). The species using the next most energetic resource (Fig. 2b, orange) then grows the fastest in the adjacent region, and similarly inhibits the growth of others (Fig. 2b, green). This process continues as inhibited species and unconsumed resources diffuse further away from the source at *x* = 0.

The resulting pattern of resource profiles is similarly stratified, as resources with progressively decreasing energy content are depleted deeper and deeper away from the source *x* = 0 (Fig. 2c). We refer to this spatial pattern as energy-ordered resource stratification.

To quantify the degree of energy-ordered stratification, we first define the penetration depth of each resource. Specifically, for each resource, we define its penetration depth as the width of the rectangle with the same height as the resource concentration at the source *x* = 0, and with an area equal to that of the resource profile (quantified numerically; pictorial representation in Fig. 3a). At the penetration depth, the area of the rectangle which does not overlap with the resource profile has the same area as the remaining resource profile (Fig. 3a; shaded area). For each simulation, we measure the “stratification order parameter” as the magnitude of the correlation between the energy content and penetration depth of all simulated resources (Fig. 3b).

**FIG. 3.**
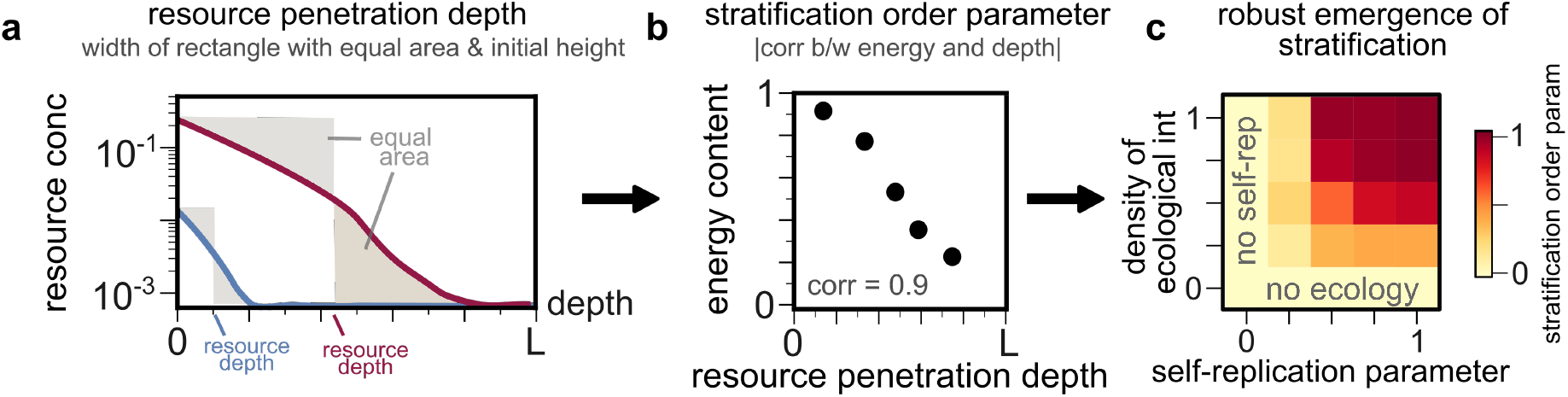
Both self-replication and antagonism are necessary for robust stratification. (a) Quantification of resource penetration depth: for each simulated resource profile (blue and red), we find the width of the rectangle with area equal to that of the resource profile and the same initial height. (b) Quantification of stratification order parameter: for all resource profiles obtained from one simulation, we compute the magnitude of the correlation between their penetration depths and energy content *Y*_*α*_ (shown is an example from a simulation with 5 profiles). (c) Heatmap of the stratification order parameter over multiple simulations, where we systematically varied the self-replication parameter *γ* and the density of ecological interactions *ρ*. Stratification does not emerge in the absence of either selfreplication or ecology. As *γ* and *ρ* both increase, stratification emerges robustly.

To test the robustness of energy-ordered stratification, we repeat 1,000 simulations of our model across a variety of randomly chosen conditions (see Methods). Throughout simulations, we systematically vary two key parameters: the self-replication parameter *γ* and the density of ecological interactions *ρ*. In each case, we quantify the mean stratification order parameter across simulations. We find that self-replication and ecology are not only sufficient, but also necessary to generate energy-ordered stratification (Fig. 3c).

In the absence of self-replication (*γ* = 0), the energy content of a resource has no bearing on its spatial profile, with penetration depth set by diffusion and consumption rate (not energy content). In the absence of ecology (*ρ* = 0), species coexist with no spatial segregation; as a result, all resources are co-utilized to depletion, with energy-ordered stratification again failing to emerge. In the presence of both, energy-ordered stratification emerges robustly, with the stratification order parameter rapidly transitioning to ≈1 as both *γ* and *ρ* increase beyond 0 (Fig. 3c; dark red region). Taken together, self-replication and ecology are both sufficient and necessary to generate self-organized stratification of resources in order of their usable energy content.

## DISCUSSION

So far, the search for chemical signatures of extraterrestrial life has been largely confined to looking for specific molecules believed to be associated with biotic processes (e.g., phosphine [18]). This approach is almost inevitably limited to forms of life closely resembling those on Earth. Until recently, such candidate biosignatures have been the only feasible approach, since our access to faraway worlds has been limited to spectroscopic analysis. However, the advent of missions allowing direct probing of Mars soil or the “sample return missions” such as Hayabusa [19] or OSIRIS-REx [20] has opened up new possibilities.

Here, we proposed a new kind of biosignature, namely observing stratified spatial profiles of chemical compounds in order of their energy content. We showed that the combined action of two key ingredients—self-replication and interspecies antagonism—naturally leads to such a self-organized stratification. This is because (1) self-replication produces an emergent correlation between the energetic yield of a resource and its rate of depletion; and (2) the interspecies antagonism then allows metabolic processes to become spatially segregated. We used a minimal model to argue that these ingredients are both sufficient and necessary. This mechanism does not depend on specific hypotheses about the chemical implementation of the organisms or their metabolism, but only on the plausible assumption that for a sufficiently evolved life form, usable chemical energy in a compound (i.e., its growth yield on this resource) correlates with its total chemical energy. Thus, we propose that energy-ordered resource stratification might serve as a robust agnostic biosignature.

Stratified structures can also emerge abiotically, e.g., sedimentation and calcification [21]. However, these abiotic mechanisms are driven by solubility or other chemical properties, and are not expected to be correlated with the total chemical energy in a compound. In contrast, biotically, such a correlation emerges naturally, as our model illustrates. While we intentionally kept the ingredients to a minimum, other biotic mechanisms, e.g. metabolic cross-feeding, could lead to similar outcomes (e.g., Winogradsky columns [15, 22]). Broadening the range of biotic mechanisms supporting energy-ordered stratification as a biosignature could potentially make our proposal even more robust.

## Acknowledgements

We are grateful to A. Murugan, R. Braakman, P. Byrne, G. Fournier and S. Seager for helpful discussions. This work was supported in part by the NSF grant PHY-2310746. A.G. acknowledges support from the Ashok and Gita Vaish Junior Researcher Award, as well as the Government of India’s DBT Ramalingaswami Fellowship.

## METHODS

We simulated our model by numerically evolving equations (1)–(2) with the assumptions as in equations (3) and (4). All simulations were done on a 1D domain of length *x* ∈ [0, *L*] where *L* = 100 in arbitrary units, assuming no boundary flux Neumann boundary conditions for species, and assuming resources entered at *x* = 0 at flux *K* and had no flux at *x* = *L*. For all simulations, we set *D*_*N*_ = 10, *D*_*R*_ = 20, *m*_*i*_ = *m* = 0.1 and *k*_*iα*_ = 1 for all species and resources. For each simulation with *M* resources and *S* = *M* species, we chose the energy content *Y*_*α*_ of each resource *α* randomly from a uniform distribution between 0. For competitive interaction strengths *α*_*ij*_ between species *i* and *j*, we first picked a fraction *ρ* of the interactions randomly, setting the rest to zero. For the picked interaction strengths, we chose them by randomly selecting a number from a normal distribution with mean 0.4 and standard deviation 0.1. All diagonal entries *α*_*ii*_ = 0 were left out of this procedure. For initial conditions, we always set homogeneous initial conditions for species, while choosing quadractically decaying profiles for resources, numerically obtained to satisfy the boundary conditions.

## Notes

### Competing Interest Statement

The authors have declared no competing interest.

